# Synergistic effects of conductivity and cell-imprinted topography of chitosan-polyaniline based scaffolds for neural differentiation of adipose-derived stem cells

**DOI:** 10.1101/2020.06.22.165779

**Authors:** Behnaz Sadat Eftekhari, Mahnaz Eskandari, Paul Janmey, Ali Samadikuchaksaraei, Mazaher Gholipurmalekabadi

## Abstract

Smart nano-environments that mimic the stem cell niche can guide cell behavior to support functional repair and regeneration of tissues. The specific microenvironment of nervous tissue is composed of several physical signaling factors, including proper topography, flexibility, and electric conductance. In this study, a cell-imprinting technique was used to obtain a hierarchical topographical conductive scaffold based on chitosan-polyaniline (PANI) hydrogels for directing the neural differentiation of rat adipose-derived stem cells (rADSCs). A chitosan-polyaniline hydrogel was synthesized, followed by characterization tests, such as Fourier transform infrared spectroscopy (FTIR), electrical conductivity, Young modulus, and contact angle measurements. A chitosan-PANI scaffold with a biomimetic topography was fabricated by molding it on a chemically fixed culture of PC12 cells. This substrate was used to test the hypothesis that the PC12 cell-imprinted chitosan-PANI hydrogel provides the required hierarchical topographical surface to induce neural differentiation. To test the importance of spatial imprinting, rADSCs were seeded on these conductive patterned substrates, and the resulting cultures were compared to those of the same cells grown on flat conductive chitosan-polyaniline, and flat pure chitosan substrates for evaluation of adhesion, cell viability, and expression of neural differentiation markers. The morphology of rADSCs grown on conductive patterned scaffolds noticeably was significantly different from that of stem cells cultivated on flat scaffolds. This difference suggests that the change in cell and nuclear shape imposed by the patterned conductive substrate leads to altered gene expression and neural differentiation of cultured cells. In summary, a conductive chitosan-polyaniline scaffold with biomimetic topography demonstrates a promising method for enhancing the neural differentiation of rADSCs for the treatment of neurodegenerative diseases.

---

The human nervous system consists of the central nervous system (CNS) and the peripheral nervous system (PNS), which play distinct and critical roles in all physiological processes, including recognition, sensory and motor functions of cells. ^1, 2^ Neurodegenerative disorders in the region of the brain and spinal cord, traumatic injuries, and stroke influence the quality life of 2 million people in the United States of America (USA) each year, and this number annually grows by an estimated 11000 cases.^3, 4^ The regeneration of injured neurons is limited under normal conditions due to two main factors: post-damage scar formation and unguided axonal regrowth. ^5^ The nerve injury activates the proliferation of glial cells, known as astrocytes in CNS and Schawnn cells (SCs) in the PNS, which create scar tissue and prevent the regrowth of severed axons by producing inhibitory molecules ^6^ and altering the extracellular environment. Due to the ineffectiveness of numerous strategies for the regeneration of neural defects, full recovery of damaged nerves remains challenging. ^7^

New studies have also demonstrated the potential of stem cells for the treatment of catastrophic diseases such as neurodegenerative diseases and cancer.^8, 9^ The stem cell environment (niche) controls the natural regeneration of damaged tissue through providing biochemical (e.g., growth factors and other soluble factors) as well as biophysical cues (e.g., shear stress, elastic modulus, stiffness, geometry, and conductivity). Due to disappointing clinical results of therapeutic approaches with growth factors alone (e.g., angiogenic factors), fabrication of engineered scaffolds with appropriate biophysical and biochemical properties have been proposed.^10–12^

Fundamental developments in tissue engineering have made strides to control stem cell behaviors such as proliferation, migration, and differentiation by applying biophysical signals ^13^. In particular, nerve tissue engineering scaffolds must present appropriate physical characteristics supplying the proper topography, mechanical elasticity, and electrical conductivity to accelerate axonal growth during regeneration.^14^ Micro and nano topographical substrates can actuate the mechano-transduction pathways that regulate stem cell fate by contact guidance through the rearrangement of cytoskeleton alignment, leading to changes in nucleus shape and altering gene expression.^15^

Most studies have indicated that nanopatterned surfaces activate cell surface proteins such as integrins, which are responsible for cell surface signal transduction, cluster assembly, and formation of focal adhesion complexes containing vinculin, paxillin, and focal adhesion kinase (FAK), which alter the arrangement and mechanical tension of the cytoskeleton.^16, 17^ Ultimately, the mechanotransduction pathway activated by mechanical tension changes the nucleus shape and the expression profile of genes involved in stem cell differentiation.^18^ Since the stem cell niche comprises nano and micropatterned topographies, scaffolds with various geometries such as continuous topography, discontinuous topography and random topography with different spatial dimensions ranging from nanometers to micrometer (which are typically fabricated by photolithography, microcontact printing, microfluidic patterning, and electrospinning techniques) are utilized for regulation of stem cell fate to neural differentiation.^19, 20^ Despite much progress to attain precise control of stem cell behavior using engineered patterned substrates, high yield, reliable, safe, and cost-effective control of stem cell fate remains a challenge.

in addition to the role of substrate topography, conductive scaffolds that mimic the electrical conductivity of native tissue promote stem cell plasticity and differentiation into specific lineages by altering their membrane depolarization.^26^ Because of the involvement of endogenous electrical signals in neurogenesis, nerve growth, and axon guidance, biophysical studies of electrical signals are increasingly used to direct stem cell differentiation toward neuron-like cells.^27^ Electrically conducting polymers such as polypyrrole, polyaniline (PANI), polythiophene, and their derivatives (mainly aniline oligomers and poly (3,4-ethylene dioxythiophene)) are bioactive biomaterials for controlled delivery of electrical signals to cells. ^28^ Despite the biocompatibility of these polymers, their weak mechanical properties and poor processability require blending these polymers with other biomaterials.^28^ Chitosan (CS), as a biocompatible, biodegradable, non-immunogenic, and antibacterial biomaterial, is frequently considered a promising candidate for electroactive hydrogel fabrication.^29–31^ Hence, we proposed to examine potential synergetic effects of cell-imprinted conductive CS-PANI hydrogels on the differentiation of rat adipose-derived stem cells (rADSCs) into neuron-like cells. In the present *in vitro* study, we synthesized pure CS and conductive CS-PANI hydrogels. In the next step, a chemically fixed differentiated culture of PC12 cell was employed as a template on which the conductive hydrogel was cast. After the removal of the remaining cells/debris, the conductive scaffold was obtained with the specific topography of the cells that were used as a template. Consequently, the potency of the cell-imprinted conductive scaffold and a control flat conductive scaffold was investigated for inducing the neural differentiation of rADSCs.

## RESULT AND DISCUSSION

### Fourier-transform infrared spectroscopy (FTIR) spectra

The FTIR spectra of pure chitosan and PANI/CS blend substrates are shown in **Figure 2**. In the pure chitosan spectrum, the quite broad peak at 3320 cm^−1^ can be assigned to the overlapping of OH and NH2 stretches. The bands occurring at 2918 cm^−1^ and 2873 cm^−1^ are ascribed to the C-H stretching of the aliphatic group. The transmission peaks at 1660 cm^−1^ belong to the C=O in amide groups (NHCOCH3) because of the partial deacetylation of CS. N-H bending was observed at 1554 and 1416 cm^−1^. The transmission peak at 1386 cm^−1^ is due to the C-OH vibration of the alcohol groups in CS. Other main peaks observed in CS spectra involving 1299 cm^−1^, 1254 cm^−1^, and 1144 cm^−1^ are ascribed to anti-symmetric stretching of the C-O-C bridge and the C-O stretching, respectively.^32, 33^ After blending PANI with CS, a small shift of peaks can be seen for PANI–CS composite with N–H stretching vibrations at 3290 cm^−1^, C–H stretching and vibrations at 2887/2861cm^−1^, respectively. A slight shift of peaks was also indicated with the amide I and II vibrations at 1646 cm^−1^ for CS-PANI composite. Because of interactions between chitosan and polyaniline and conformational changes, small shifts occurred on the C–H bending vibration of the amide methyl group (1386 cm^−1^) and the C–O stretching vibrations at 1161-1034 cm^−1^. The characteristic transmission bands of PANI were also authenticated in the CS-PANI blend spectrum. The transmission peaks corresponding to the C=N stretching vibration of the quinonoid ring and the C=C stretching vibration of the benzenoid ring can be seen at 1554/1519 cm^−1^, respectively.^34, 35^ The presence of these peaks confirms that the synthesized composite samples contained PANI.

**Figure 1:**
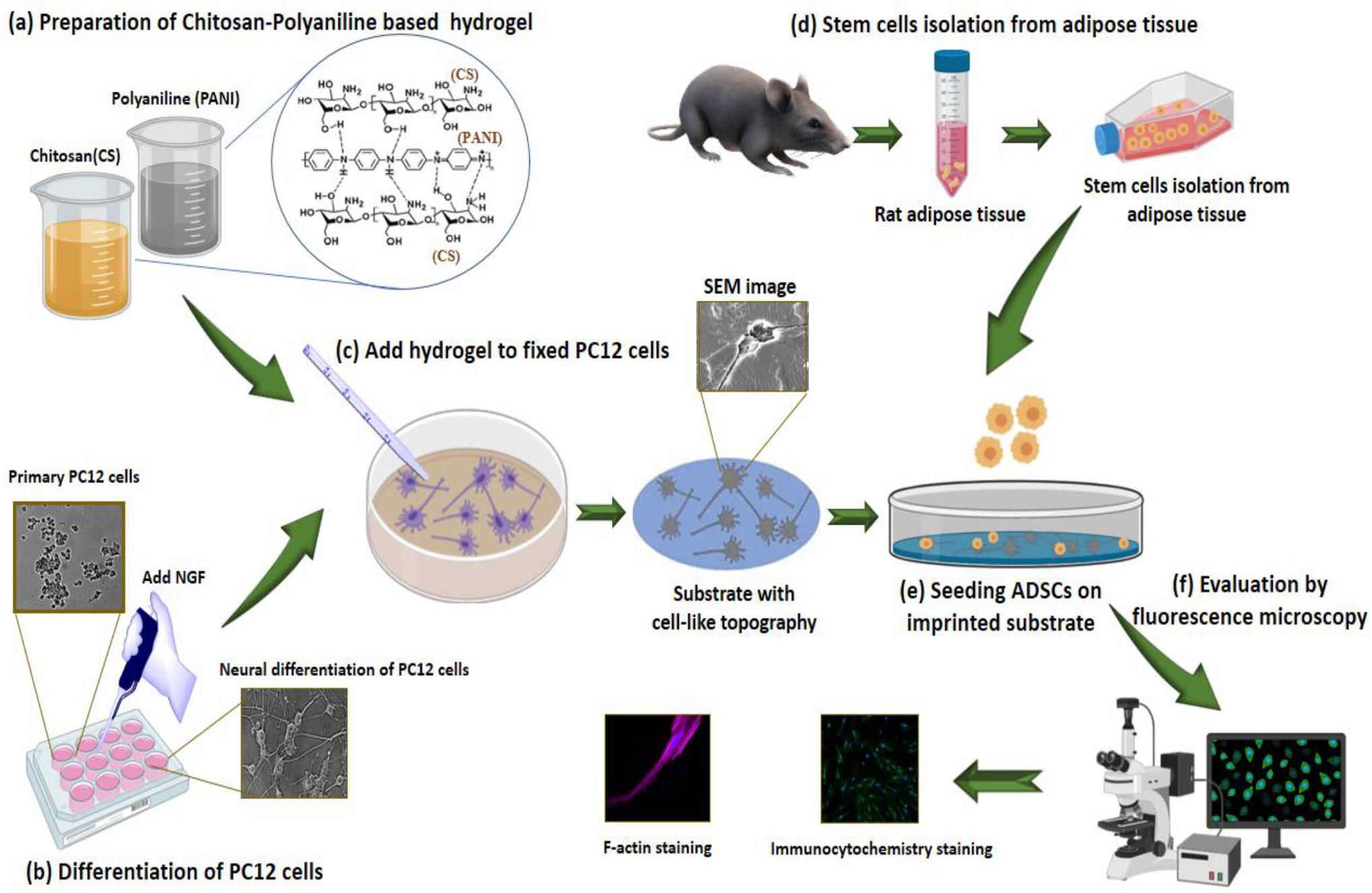
Schematic represents the experiment steps: (a) Chitosan-polyaniline based hydrogel was prepared, and (b) PC12 cells were differentiated into neural cells using NGF. (c) After the fixation of these cells, PC12 morphologies were transferred to prepared hydrogel by mold casting. In the next step, (d) stem cells were isolated from rat adipose tissue, and (e) these cells were cultured on the imprinted substrate. After 8 days, (f) the neural differentiation of rADSCs was evaluated.

**Figure 2:**
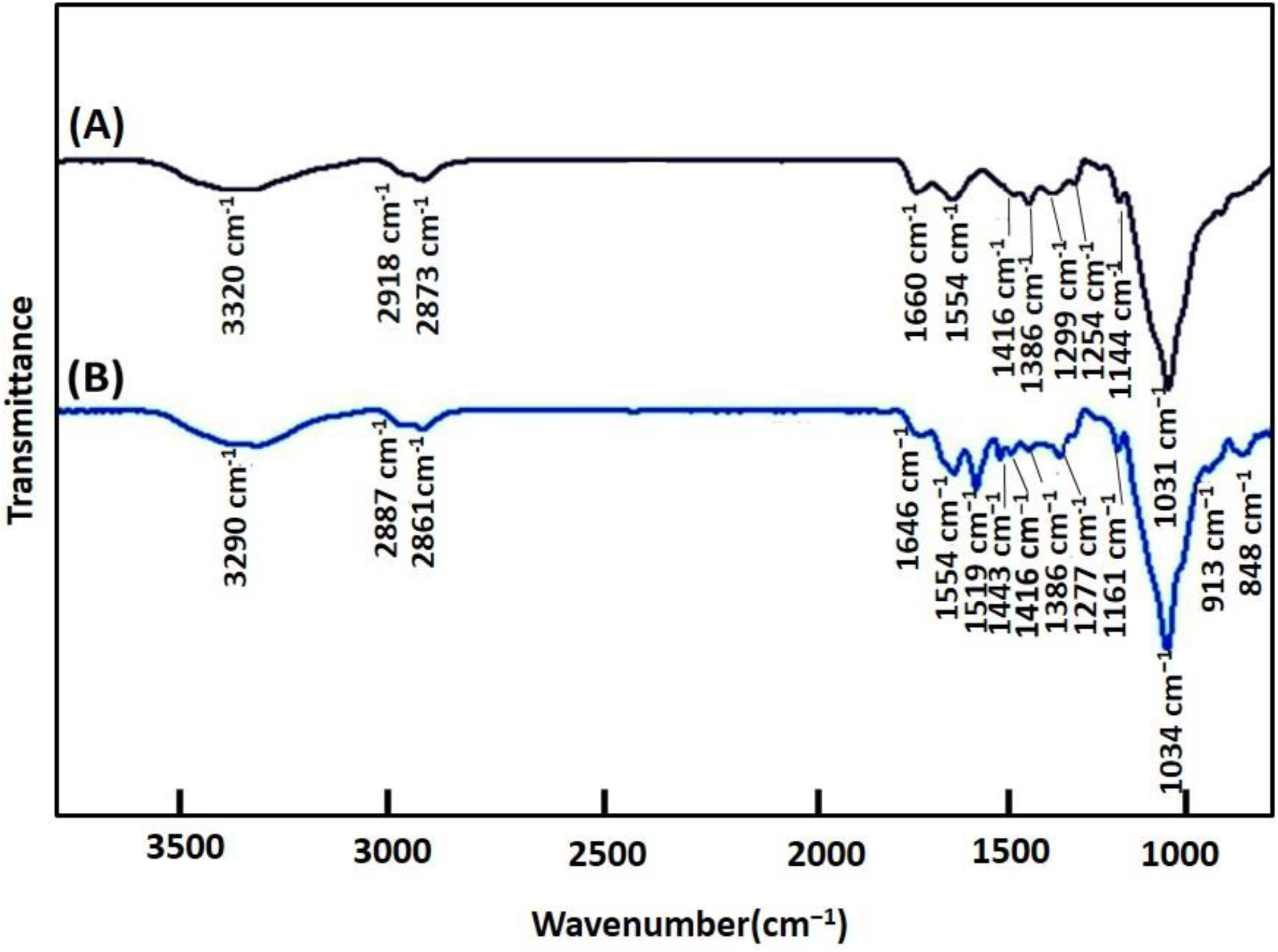
FTIR spectra of A) CS and B) CS-PANI blend substrate are shown. The exhibition of peaks at 1646 cm^−1^, 1519 cm-^1^, 1443 cm^−1^, 1277 cm^−1^, and 1144 cm^−1^ peaks in the spectra of CS-PANI blend confirm that this substrate includes PANI.

### Electrical conductivity

**Figure 3(a)** illustrates the electrical conductivity of all prepared substrates. The conductivity of the substrates was increased from 7.5×10^−8^ to 10^−4^ S/cm by enhancing the amount of PANI from 0 to 2.5 wt. % (P ≤ 0.0001). The pure chitosan substrates have weak conductivity 7.5×10^−8^ S/cm because of the polarity of protonation of the amino group. We obtained similar results for both flat and imprinted CS substrates. Hence, in the present study, we use a conductive polymer such as PANI for blending with CS to enhance the electrical conductivity of the flat and patterned substrates. The highly π-conjugated system of PANI strongly affected the electrical conductivity of the blend substrate.^34^ The results show 4 orders of the magnitude increase in specific conductivity, and there was no meaningful difference between the flat and imprinted substrates. These ranges of conductivity are sufficient for the conduction of electrical signals in *in vivo* condition [25].

**Figure 3.**
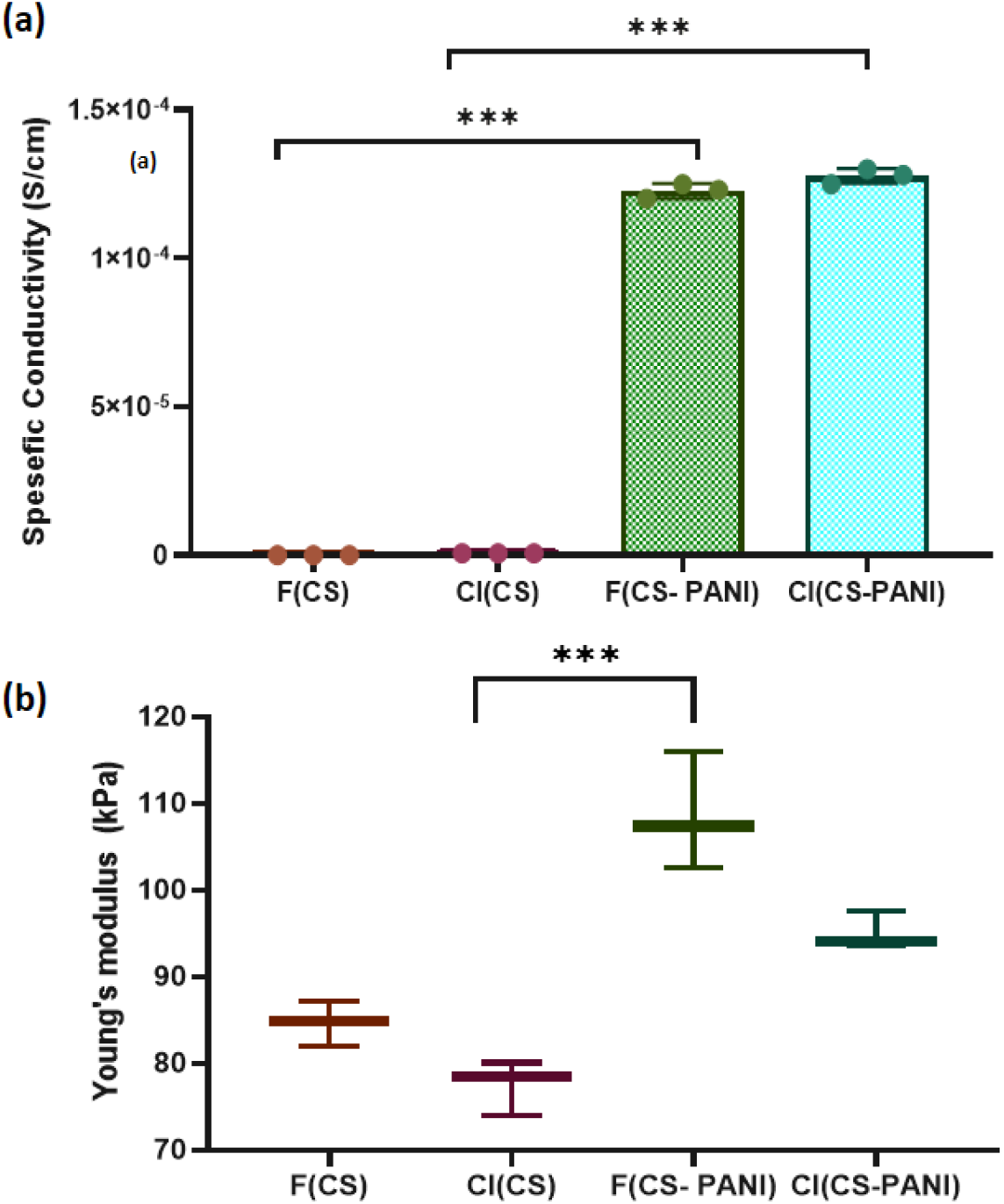
(a): The electrical conductivity of the substrates was increased by adding PANI to pure CS.

### Mechanical properties

According to the mechanical properties of stem cells, the niche can regulate cell behavior such as attachment, migration, and differentiation, so these physical cues have been considered as an essential factor in designing the artificial microenvironment to direct the cell fate.^41^ Thus, the scaffold used for nerve tissue engineering must mimic the mechanical properties of the ECM to promote the neural differentiation of stem cells.^42^ The elastic modulus (Young’s modulus) of substrates at different blend compositions are shown in **Figure 3(b)**.

### Contact angle

The hydrophilicity of the substrate is an important characteristic that affects cell attachment. Hence, the surface wettability of the flat and cell-imprinted substrates was measured by water contact angle after treatment with NaOH solution **(Figure 4(a))**. Before neutralization, the hydrophilic nature of CS-PANI blends causes the water droplet to absorb too rapidly for the contact angle to be measured. The contact angle increases between 40°-60° by adding PANI to the substrates. Also, the cell-imprinted CS and CS-PANI substrates were slightly more hydrophobic compared to flat CS and CS-PANI, respectively.

**Figure 4.**
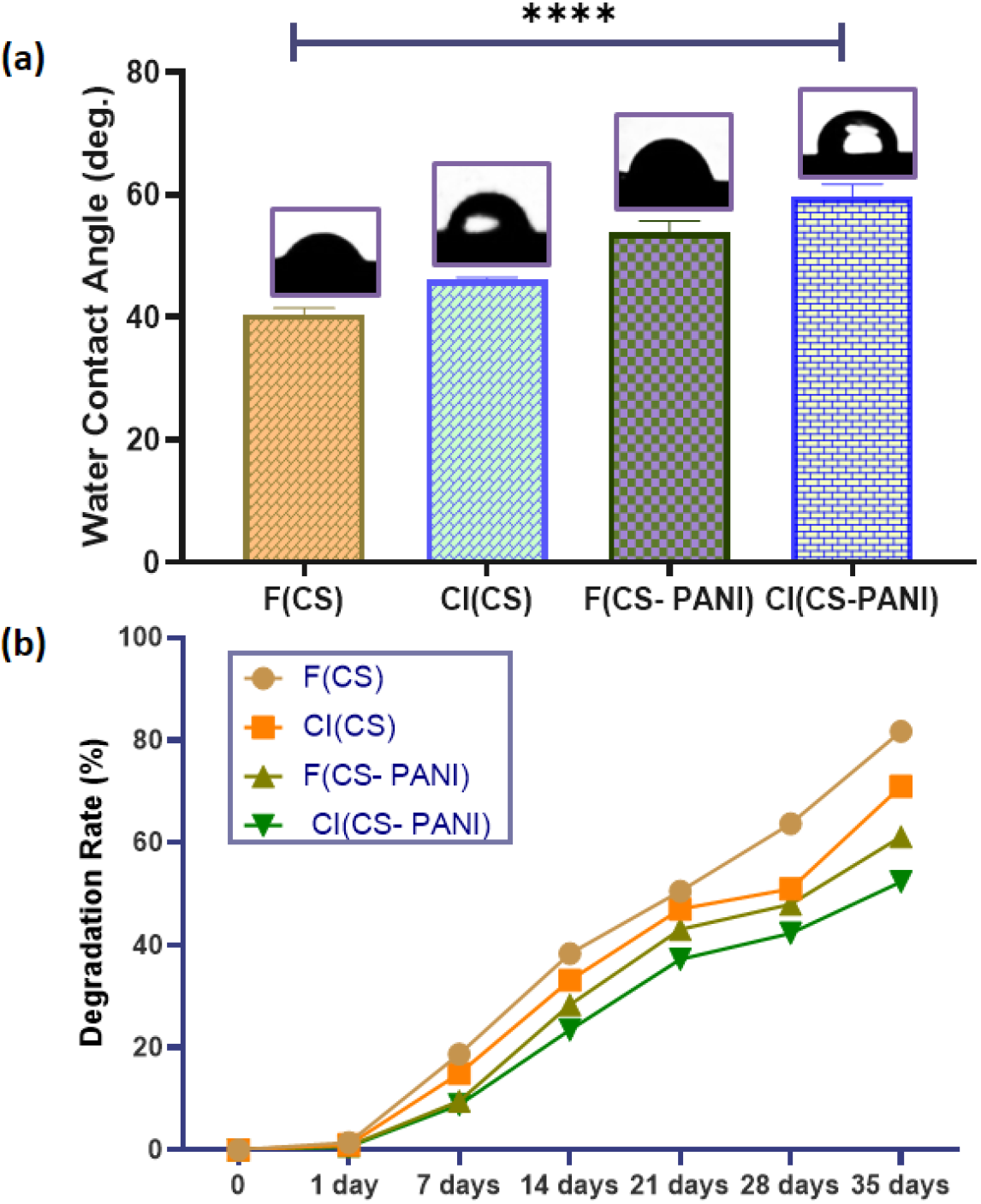
(a): Characterization of wettability of various samples. Water contact angle measurement. (b): *In vitro* biodegradation of prepared substrates in PBS was examined over 5 weeks. ****P < 0.0001.

### Degradation assay

Throughout *in vivo* tissue regeneration, the substrate should provide the support structure for cell attachment, proliferation, and differentiation. Successful regeneration for peripheral nerve damage takes 5-8 weeks.^47^ Since one of our aims was to evaluate if surface topography might alter the effect of a CS-PANI blend on neural differentiation due to altered biodegradation, the *in vitro* degradation of substrates in PBS solution was examined for 5 weeks. **Figure 4(b)** shows that the degradation rate of pure CS substrates was faster than other groups, but still preserved approximately 60% of its initial mass over 2 weeks.

### Characterization of rat adipose-derived stem cells (rADSCs)

Fibroblast-like and specific CD markers on the cell surface are the most important criteria for the characterization of ADSCs.^48, 49^ These cells are positive for mesenchymal specific markers such as CD29 and CD90 and negative for hematopoietic stem cell markers such as CD34 and CD45.^50^ The morphology of ADSCs after 5 and 14 days after isolation is shown in **Figure 5(a)**. These cells displayed the adherent and typical fibroblast-like morphology under optical microscopy. Flow cytometry results demonstrated that 99.4% and 98.85% of the cell population were positive for CD90 and CD29, while only 0.122% and 0.457% of them were positive for CD34 and CD45, respectively **(Figure 5(b))**. Immunostaining by CD surface markers was also performed (**Figure 5 (c-f))**. These results revealed that more than 85% of rADSCs positively expressed the mesenchymal stem cell markers CD29 and CD90 but did not express the hematopoietic stem cell marker CD45 and CD34 (< 5%).

**Figure 5.**
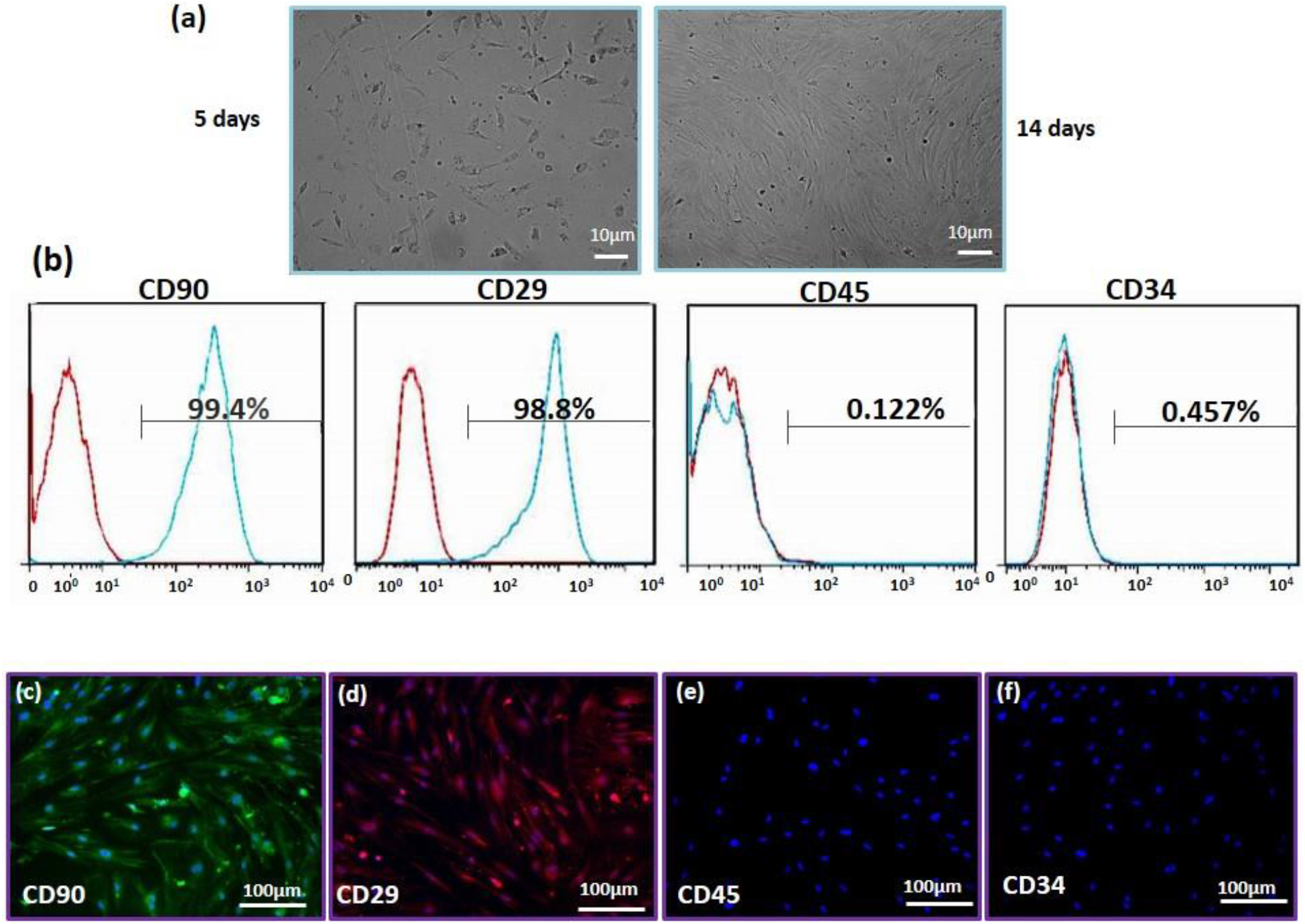
(a): Morphological change in ADSCs in cell culture media at days 5 and 14 post-isolation. (b): Flow cytometry results of CD90, CD29, CD45 and CD34. More than 90% of cell population represented phenotypic characteristics of ADSCs. Immunostaining of the rADSCs expression of (c):CD90 (green), (d): CD29 (red), (e): CD45 and (f): CD34. Cell nuclei were stained with DAPI (blue). The result of staining confirmed the flow cytometry results.

### Cell viability study(MTT assay)

The biocompatibility of pure CS substrates and CS-PANI substrates was evaluated by MTT assay to define the effect of the PANI concentration on rADSCs proliferation on 1, 3,7,14 days after cells seeding. The results from the determination of proliferation rate **(Figure 6)** showed no significant difference between cells cultured on typical TCP and cells cultivated on pure CS substrates or CS-PANI substrates. The MTT results confirm the biocompatibility and supporting role of flat and cell-imprinted CS-PANI substrates for adhesion and growth of ADSCs. This results is important, considering that conductive polymers such as PANI must be used in optimal concentration to be compatible for cells.^54^

**Figure 6:**
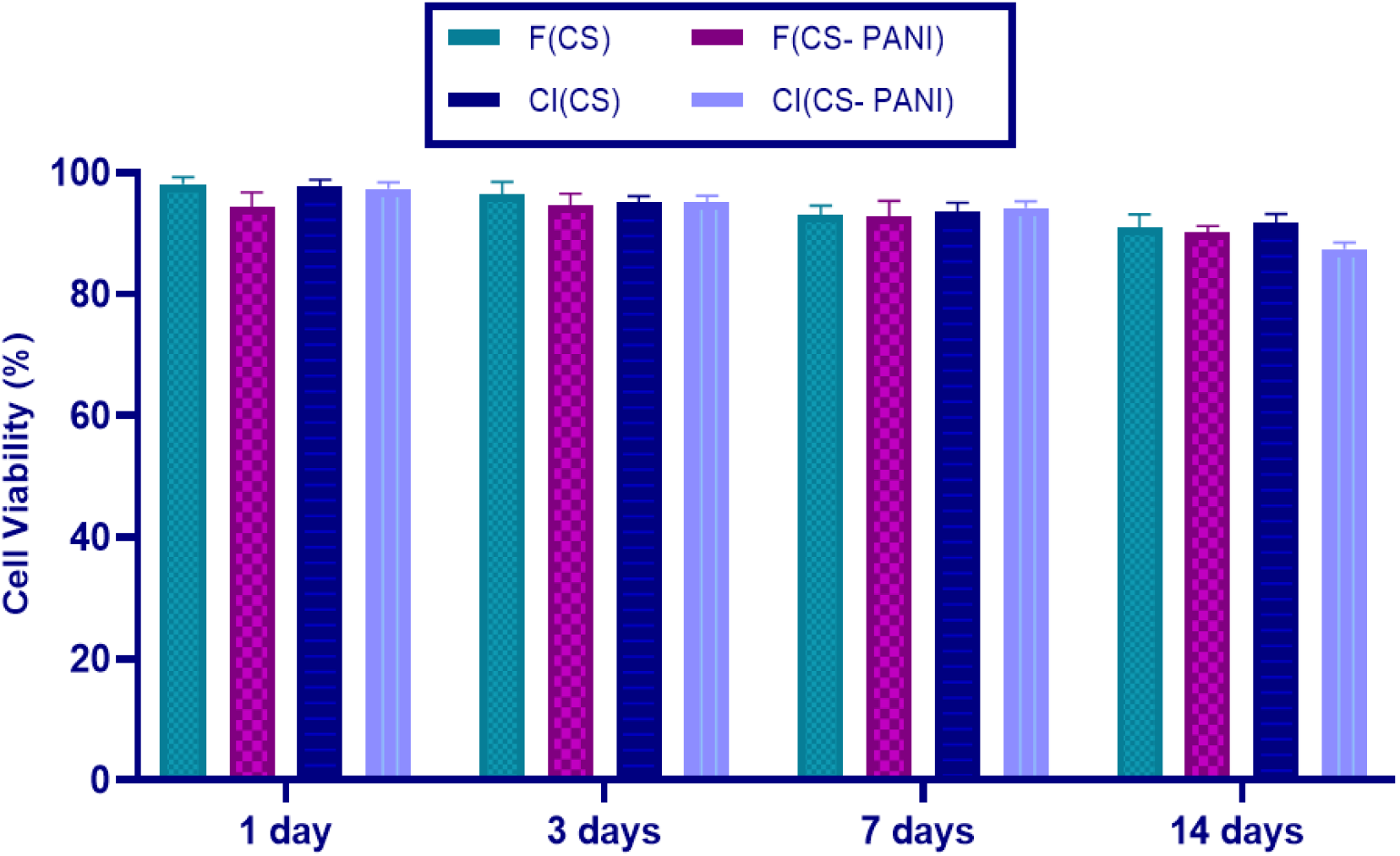
MTT viability assay of cultured ADSCs on the prepared substrates. F (CS): Flat chitosan substrate, F (CS-PANI): Flat chitosan-polyaniline substrate, CI (CS): cell-imprinted chitosan substrate, CI (CS-PANI): cell-imprinted chitosan-polyaniline substrate. (* P < 0.001).

### Immunocytochemistry

The effect of cell-imprinting topography and conductivity of the scaffolds on neural differentiation of ADSCs was examined by staining them with MAP2 and GFAP specific antibodies and viewed by fluorescent microscopy (**Figure 7**). MAP2 isoforms are expressed only in neuronal cells, while GFAP is expressed in astrocyte cells and contributes to many important CNS processes, including cell communication and the functioning of the blood brain barrier.^55^ After 8 days, more ADSCs on imprinted substrates were more immunoreactive for GFAP and MAP2 markers relative to ADSCs that, were cultured on flat substrates, suggesting significant the topography-driven neural differentiation. Furthermore, by adding PANI to the substrate, the level of expression of neural-specific genes and, consequently, the degree of differentiation was enhanced.

**Figure 7.**
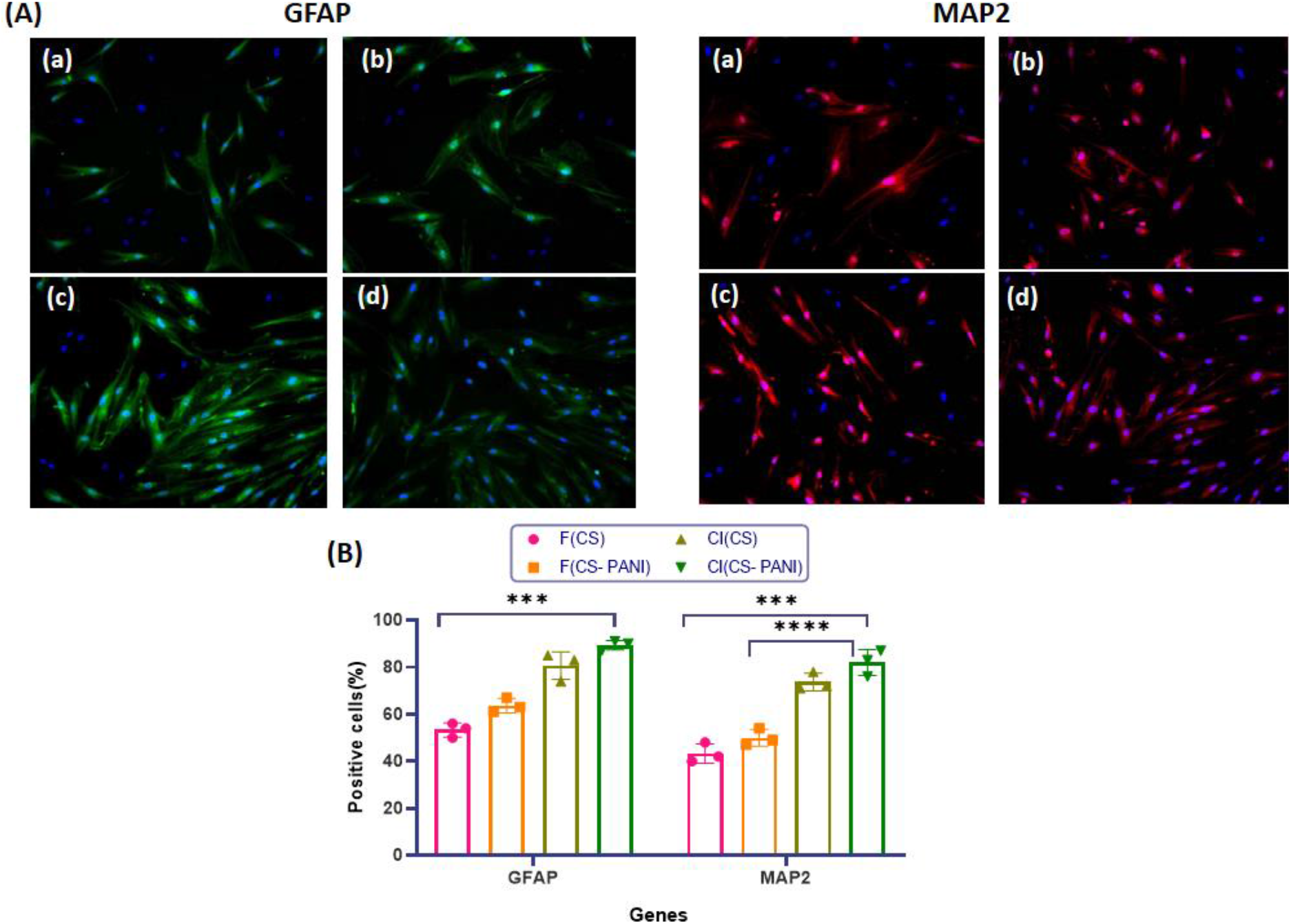
(A) Immunostaining of the rat adipose derived stem cell (rADSCs) expression of GFAP (green) and MAP2 (red) markers. Cell nuclei were stained with DAPI (blue). Scale bar: 100 μm. (B) Average percentage of GFAP and MAP2 expressing rADSCs. (a): F(CS); (b): F(CS-PANI); (c): CI(CS); (d): CI(CS-PANI). p ≤ 0.05 was considered as level of significance. * indicates significant difference.

## CONCLUTIONS

The substrates produced for neurogenic differentiation effectively simulate the natural ECM of neuron cells in multiple aspects. In this study, conductive cell-imprinted substrates mimic the topography and conductivity of nerve tissue, which can physically direct stem cells toward neuron like cells. PC12-imprinted CS-PANI substrates exhibit desirable characteristics such as appropriate mechanical properties, degradation rate, and good wettability, which represent effective parameters stimulating neural differentiation. Finally, this work prepares a pioneered design of patterned conductive scaffolds for controlling the differentiation of ADSCs, which has the potential for effective stem cell therapy of neurological diseases and injury. This synergetic effects and the comparison between the mechanical strength, topography, and electro-conductivity propose that by trying to bring more cues to the neural tissue engineering, mechanotransduction can be used as one of the main reasons for nonconventional results but at the main time push us toward thermodynamic parameters of the cell structure.

## EXPERIMENTAL SECTION

### Materials

Polyaniline (PANI, emeraldine base, Mw 65,000), chitosan (%DD=80, medium molecular weight), and 3-(4,5-dimethylthiazol-2-yl)-2,5-diphenyl tetrazolium bromide (MTT) were purchased from Sigma-Aldrich. Glutaraldehyde (50% w/w, analytical grade) and paraformaldehyde were acquired from Fluka. AR grade chemicals of N-methyl-2-pyrrolidone (NMP), acetic acid, and methanol were used as received.

### Fourier-transform infrared spectroscopy (FTIR) spectra

A Thermo Nicolet Nexus FTIR spectrometer in the transmittance mode at 32 scans with a resolution of 4 cmK was used for recording the FTIR spectra of chitosan and the blend samples. Spectra in the frequency range of 4000– 400 cm-^1^ were measured using a deuterated tri-glycerine sulfate detector (DTGS) with a specific detectivity of 1*10^9^ cm Hz1/2 wK1.

### Imaging

The surface morphology of the topographical substrates and flat substrates was investigated by scanning electron microscopy (SEM) (Hitachi Japan; apparatus working at 10 keV accelerating voltage), atomic force microscopy (AFM; DME DS 95Navigator 220) and optical microscopy (Nikon Optiphot 200). Before SEM imaging, all samples were coated with a thin layer of gold using a sputtering machine. AFM contact mode was performed using a rectangular cantilever (HQ:NSC18/Al BS, MikroMasch, Bulgaria) with a spring constant of 2.8 N/m and a conical tip of 8 nm radius. The surface of the sample up to 90 μm^2^ was scanned by a scan rate of 0.05 Hz and setpoint force 0.5 nN. The standard software of the instrument (JPKSPM Data Processing) was utilized for image analysis.

### Electrical conductivity

The electrical conductivity of fabricated substrates was measured by applying a two-point probe method (Keithley, model 7517A). The electrical conductivity (σ) is obtained as the inverse of resistivity. The resistivity of three samples from each group of the substrate was measured by passing a constant current through the outer probes and recording the voltage via the inner probes. The resistivity of samples was calculated as follows:

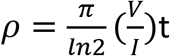

where: I, V, and t indicate the applied current, voltage, and the sample thickness (200 μm), respectively. ^57^

### Mechanical test

The substrates were immersed in PBS to reach the equilibrium swelling before performing mechanical tests. The mechanical properties of all fabricated samples were determined using AFM. A Molecular Imaging Agilent Pico Plus AFM system (now known as AFM5500 Keysight technologies) with silicon nitride probes and 5 μm spherical (nominal value) tips (CP-PNPL-BSG, and a spring constant of 0.08Nm^−1^ was used to measure the Young’s modulus of samples.

### Contact angle

The static (sessile drop) water contact angle of the prepared substrates was assessed using a contact angle apparatus (DSA20, Germany) at room temperature. A droplet of ultra-pure water was placed on the fixed sample surface, and the measurement was performed 3 s after equilibration. Then, a camera recorded the water contact angle of each surface. This measurement was repeated at three or more different locations on each sample to calculate the average value of the contact angle.

### Degradation rate

The *in vitro* degradation of substrates was followed for 5 weeks in PBS (pH =7.4) at 37 °C. Three samples from each group with ~40 mg weight were immersed in 10 ml of PBS at pH = 7.4 and 37°C. At time points of 3, 7, 10, 14, 21, and 30 days, each of the samples was taken out and dried for 12h after surface wiping, and its weight was recorded. The following equation was used for the calculation of sample weight loss:

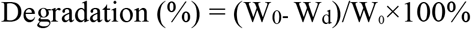

Where W_0_ is the dry weight of the sample at time t = 0 and W_d_ is the dry sample weight after removal from the solution. The pH values of the solutions during scaffold immersion were also recorded.

### Isolation and Culture of rat Adipose-derived Stem Cells (rADSCs)

Rat adipose-derived stem cells (rADSCs) were isolated and expanded by a procedure described in a published study. ^58^ In brief, subcutaneous adipose tissue samples were isolated by needle-biopsy aspiration according to a previously published methodology. DMEM medium (GIBCO, Scotland) containing 10% (v/v) FBS and penicillin (100 IU/ml)-streptomycin (100 μg/ml) was used for transportation to the cell culture laboratory. After washing the tissue sample with DMEM-based buffer three times, the samples were gently cut into small pieces and incubated with 0.05 mg/ml collagenase type I (Sigma, USA) for one 1h to digest the epididymal fat. The separated cells were collected from the resulting suspension after centrifugation at 200 ×g for 5 minutes. The rADSCs were resuspended in DMEM supplemented with 10% FBS, 100 U/mL penicillin and, 100μg/mL streptomycin in an incubator (37°C, 5% CO2). The culture medium was changed to remove non-adhered cells and debris.

### Characterization of rADSCs by flow cytometry and immunocytochemistry

The rADSCs were examined by flow cytometry for identification of surface markers. The cells were identified for expression of mesenchymal markers such as CD29 and CD90 and the hematopoietic markers CD34 and CD45 (all antibodies obtained from Novus Biologicals company, US). Briefly, 1×10^6^ cells in PBS were incubated with 1 μg of each antibody for 1 hour and then washed with PBS for 3 times. Ultimately, cells were treated with secondary antibodies and protected from light for 30 minutes. All cell preparations were analyzed by flow cytometry (FACSCalibur, BD), and data analysis was performed with FlowJo software (Tree Star). The rADSCs were also characterized by immunocytochemical staining as follows: the cells were fixed with 4% PFA in PBS for 15 min and permeabilized with 0.25% Triton X-100 in PBS for 10 min at RT. 3% BSA as blocking solution was added to samples for 30 min and incubated with the primary antibodies (diluted block solution) overnight at 4 °C. Fluorochrome-conjugated secondary antibodies were then added to samples for 1 h at RT and DAPI solution for another 10 min to stain the nucleus. Finally, the sample was rinsed with PBS 3 times. Fluorescence signals were detected using a Leica DMIRE2 microscope under the proper exciting wavelength.

### Stem Cell Seeding on cell-imprinted substrates

The prepared substrates were placed into the 6-well plates. rADSCs (3×10^3^ cells/cm^2^) were then cultured on the conductive cell-imprinted substrates, conductive flat substrates, and flat pure chitosan films. After 24 h, 600 μL/cm^2^ of fresh DMEM with 10% (v/v) FBS was added to each well to cover the substrates completely.

### Cell Viability

The viability of rADSCs cells was investigated by an MTT assay. To perform this test, a cell density of 2 × 10^4^ cells/cm^2^ were cultured on flat and cell-imprinted CS and CS-PANI substrates for 1 day, 3 days, 7 days and 14 days in the incubator (37°C, 5% CO2). At these specified times, the cells were washed with PBS, and then the MTT solution was added into well. After incubation for 4 h, live cells created formazan crystals and were washed with PBS. ADMSO/isopropanol solution was added to dissolve the crystals. Finally, the Optical Density (OD) was measured using a spectrophotometer at a wavelength of 570 nm. The following formula determined the absorbance value:

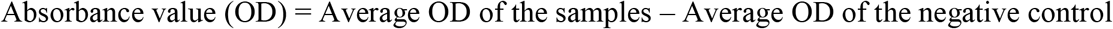

All data of the test were expressed as means ± standard deviation (SD) for n = 4. The Kolmogorov–Smirnov test was applied to investigate the normal distribution of each group of samples. Finally, a one-way ANOVA and Duncan test (GraphPad Prism 7 software) was performed to compare the data. P<0.05 was considered a significant difference.

### Neural lineage induction

The rADSCs of the 4th passage were seeded on a six-well culture plate at a density of 5 ×10^5^ cell/ml with 1 ml of neurosphere medium containing serum-free DMEM/F12 with 2 % B27 supplement (Thermo Fisher, USA) and 20 ng/ml of basic fibroblast growth factor (bFGF) (Thermo Fisher, USA) for 3 days. The neurospheres were seeded on the 6-well plate and cultured in the neurosphere medium containing 5% FBS. After 2 days, the cells were trypsinized by trypsin/EDTA (0.05 % trypsin/ 0.5 mM EDTA) and seeded on substrates in a 24-well plate. The cell suspension containing 1 × 10^5^ cell/ml density was added to the top of the plate. The 3D culture was promoted in the neurosphere medium containing retinoic acid (0.1 M) for five days.

### Immunostaining

The cell morphology and expression of neuron cell-specific markers were investigated using optical and fluorescence microscopy (Nikon, Japan) to evaluate the neural differentiation capability of rADSCs on prepared substrates. After 8 days from neural induction, the cells seeded on substrates were rinsed with PBS and fixed with 4% PFA in PBS (pH 7.4) for 20 min. In the next step, the cells were permeabilized in a solution containing 0.5% Triton X-100 in PBS for 10 min. To block non-specific antibodies, the cells were incubated with 10% normal goat serum for 1 h at room temperature (0.05% Tween 20 and 1% (w/v) bovine serum albumin (BSA)). Detection and characterization of neuronal cells were carried out by staining the cells with antibodies against GFAP and MAP2. Antibodies to GFAP (1:100; Abcam) and MAP 2 (1:50; Millipore) were added to the cell medium for staining overnight at 4°C. After cells were washed with PBS, the secondary antibody, including Alexa 488-conjugated goat anti-mouse and anti-rabbit antibodies, was added to cells, and incubated for 1h. Finally, the nuclei of cells were counterstained with DAPI. Fluorescence images were taken by an Olympus BX51 fluorescence microscope.

### Statistical analysis

Statistical analyses were performed using SPSS. Data are expressed as the mean – SD, and p < 0.05 was considered statistically significant.

## ACKNOWLEDGMENTS

The authors would like to thank support from Amirkabir University and Technology, NSF DMR-1720530 and CMMI-154857 US National Science Foundation.

